# A Systems Serology Approach to the Investigation of Infection-Induced Antibody Responses and Protection in Trachoma

**DOI:** 10.1101/2023.03.01.530575

**Authors:** Amber Barton, Ida Rosenkrands, Harry Pickering, Nkoyo Faal, Anna Harte, Hassan Joof, Pateh Makalo, Manon Ragonnet, Anja Weinreich Olsen, Robin L. Bailey, David CW Mabey, Frank Follmann, Jes Dietrich, Martin J Holland

## Abstract

**Background:** Ocular infections with *Chlamydia trachomatis* serovars A-C cause the neglected tropical disease trachoma. As infection does not confer complete immunity, repeated infections are common, leading to long term sequelae such as scarring and blindness. Here we apply a systems serology approach to investigate whether systemic antibody features are associated with susceptibility to infection.

**Methods:** Sera from children in five trachoma endemic villages in The Gambia were assayed for 23 antibody features: IgG responses towards two *Chlamydia trachomatis* antigens and three serovars (elementary bodies and major outer membrane protein MOMP, serovars A-C), IgG responses towards five MOMP peptides (serovars A-C), neutralization and antibody-dependent phagocytosis. Participants were considered resistant if they subsequently developed infection only when over 70% of other children in the same compound were infected.

**Results:** The antibody features assayed were not associated with resistance to infection (false discovery rate < 0.05). Anti-MOMP SvA IgG and neutralization titer were higher in susceptible individuals (p < 0.05 before multiple testing adjustment). Classification using partial least squares performed only slightly better than chance in distinguishing between susceptible and resistant participants based on systemic antibody profile (specificity 71%, sensitivity 36%).

**Conclusions:** Systemic infection-induced IgG and functional antibody responses do not appear to be protective against subsequent infection. This may be due to confounding factors increasing both past and future exposure to *C. trachomatis*, or antibody-dependent enhancement. Ocular responses, IgA, avidity or cell-mediated responses may play a greater role in protective immunity than systemic IgG.

## 1. Introduction

Obligate intracellular bacterium *Chlamydia trachomatis* is a major human pathogen with a unique biphasic lifecycle, alternating between intracellular reticulate bodies and extracellular elementary bodies (EBs). During the EB phase of its life cycle, 61% of the *C. trachomatis* outer membrane consists of major outer membrane protein (MOMP) by mass (Caldwell et al., 1981), a transmembrane porin and target for surface-binding antibodies. MOMP consists of five constant and four variable domains, of which the surface-exposed variable domains define the *C. trachomatis* serovar (Yuan *et al*., 1989). Whereas serovars (Sv) A-C cause the childhood ocular disease trachoma, D-K cause sexually-transmitted urogenital disease, and L1-L3 lymphogranuloma venereum (Faris et al., 2019). In the 1960s, several whole-organism vaccines were found to confer short-term serovar-specific immunity to trachoma (Mabey *et al*., 2014), and MOMP-based subunit vaccines hold great promise as another tool in moving past elimination towards eradication. However no vaccines against *C. trachomatis* have yet been licensed.

While cell-mediated Th1 responses are thought to be important for protection against trachoma (Hu *et al*., 2013), the role of humoral responses in protection is not well established. In human challenge studies, primary infection with ocular *C. trachomatis* conferred only incomplete, serovar-specific immunity (Dawson *et al*., 1966). Repeated infections are therefore common, resulting in conjunctival inflammation in the short-term (“active trachoma”), and scarring and trichiasis in the long-term (Hu *et al*., 2013).

The relationship between trachoma and humoral immunity in field studies is complex. Firstly, systemic antibody responses detected in the serum may have a different effect to local responses detected in the tears. For example, in one study serum anti-EB IgG was higher in controls than in those with active trachoma (Ghaem-Maghami *et al*., 1997), suggesting a protective association, while in another tear anti-EB IgG was a risk factor for active trachoma (Bailey *et al*., 1993). Antibody isotype also plays a role: anti-EB IgG is higher in those with trachomatous scarring, while anti-EB IgA is lower (Holland *et al*., 1993). No significant differences in MOMP-specific serum IgA or IgG have been observed between children with or without active trachoma, or between adults with or without trachomatous scarring (Ghaem-Maghami *et al*., 1997). Likewise, no significant differences in MOMP-specific tear IgA were found between clinical groups in Nepal (Hessel *et al*., 2001).

Given the previous conflicting relationships between humoral responses and trachoma, we hypothesized that antibody functional characteristics or antigen specificity may be important for protection against infection. We therefore used a data-driven systems serology approach (Arnold and Chung, 2018) with the aim of finding individual antibody features or overall antibody profiles associated with resistance to ocular *C. trachomatis*. This strategy has previously been successful in identifying multivariate signatures and new features of protective responses to typhoid vaccination (Jin *et al*., 2020), and distinguishing those with controlled latent tuberculosis from those with active tuberculosis (Lu *et al*., 2016). Here we apply the systems serology approach to a longitudinal study of ocular *C. trachomatis* infection. We find that IgG responses to antigens from different serovars correlate strongly with one another, despite serovar B predominating in the local area. We then investigate the association between individual antibody features and susceptibility, and assess whether susceptible and resistant participants have distinct serum antibody profiles.

## 2. Materials and Methods

### 2.1 Ethical Approval

The study design and procedures were approved by the joint Gambian Government-Medical Research Council Ethics Committee and the Ethics Committee of the London School of Hygiene & Tropical Medicine. Informed consent was obtained from the guardians of study participants on their behalf.

### 2.2 Study Recruitment and Sample Collection

Nine villages in the Gambia were selected after trachoma rapid assessment screening found active trachoma in > 20% of school age-children. 345 children aged 4-15 years from 31 family compounds were visited at baseline and fortnightly for 28 weeks (Faal *et al*., 2006). At each visit conjunctival swabs were collected in RNAlater (Ambion Europe Ltd, Huntingdon, UK), and stored at −20°C (Faal et al., 2005). DNA was extracted from swabs by one of two methods: either by QIAamp DNA Mini Kit, as previously described (Pickering *et al*., 2018), or by Roche CT/NG amplicor kit followed by concentration of positive samples using a QIAamp DNA Mini Kit (Andreasen *et al*., 2008).

160 venous blood samples (10ml) were collected at baseline in BD Vacutainer Lithium Heparin Tubes (Holland *et al*., 2006). Samples were transported to MRC laboratories within four hours, in a sealed box at ambient temperature. Plasma was separated by density gradient centrifugation using Lymphoprep (Axis-Shield Ltd, Kimbolton, UK) and stored at −30°C. Serological assays were carried out on a subset of 93 samples.

### 2.3 OmpA Serovar

OmpA serovar was determined by ompA sanger sequencing or *C. trachomatis* whole gene sequencing. Prior to sanger sequencing, conjunctival swab DNA samples underwent either one round of PCR or two rounds of PCR via a nested PCR approach, using a Veriti Thermocycler (Thermo Fisher) and following the procedure outlined in Andreasen et al. (Andreasen *et al*., 2008). PCR products were sequenced at Macrogen or Source Bioscience and were compared against all available *C. trachomatis*_sequences available in NCBI using BLAST to determine serovar.

Whole genome sequencing was performed as previously described (Last *et al*., 2018) and utilized the SureSelectXT Low Input kit. Processing and analysis of sequenced reads was performed as previously described (Pickering *et al*., 2022). Raw reads were trimmed and filtered using Trimmomatic. Filtered reads were aligned to a reference genome (A/Har13) with Bowtie2, and variants were called with SAMtools/BCFtools. Multiple genome and plasmid alignments were generated using progressive Mauve, and multiple gene alignments were generated using MUSCLE. Complete sequences of ompA were obtained using a reference-based assembly method also previously described (Pickering *et al*., 2022). Serovar was assigned using maximum blastn homology against all published *C. trachomatis* sequences.

### 2.4 *C. trachomatis* Culture

For EB IgG quantification by area under dilution curve, elementary bodies from *C. trachomatis* SvA (clinical strain A/2497), SvB (Tunis-864) and SvC (C/TW-3) were prepared by infecting Hep-2 cells at an MOI of 0.5-1 in the presence of MEM medium with 10% with foetal calf serum (FCS). Cells were centrifuged at 1800rpm for one hour at 37°C, then incubated at 37°C and 5% CO2 for two hours. Medium was then replaced by MEM with 10% FCS, 1 μg/ml cycloheximide, 0.5% glucose and 10 μg/ml gentamycin. EBs were purified by washing cells with 5ml HBSS, incubating with 0.05% trypsin/0.02% EDTA in PBS, then resuspending cells in culture medium. Resuspended cells were centrifuged at 3750rpm for 10 minutes, then resuspended in sucrose-phosphate medium (68.5g/L sucrose, 2.07g/L Dipotassium hydrogen phosphate, 1.1g/L Potassium hydrogen phosphate, 5% FCS, 0.5% phenol red, 0.05g/L Streptomycin sulphate, 0.1g/L Vancomycin hydrochloride, and 625μg/L Amphotericin B) on ice. Cells were sonicated, then EBs purified by ultracentrifugation on a 20% urografin gradient.

For neutralisation and phagocytosis assays, EBs from *C. trachomatis* SvB (Tunis-864) were prepared by infecting HeLa-229 cells (ATCC CCL-2.1, RRID:CVCL_1276) in the presence of RPMI 1640 medium (Gibco, Australia) containing 5% heat-inactivated FCS and 50 μg/ml gentamycin (Gibco, Australia) as previously described (Olsen *et al*., 2015). Briefly, after harvesting the bacteria by two high-speed centrifugation steps, the suspended bacteria were sonicated, further purified on renografin cushion by ultracentrifugation, then resuspended in 250 mmol/L sucrose, 10 mmol/L NaH2PO4, 5 mmol/L l-glutamic acid (SPG buffer) and stored at - 80°C. The inclusion-forming units (IFUs) of the batches were quantified by titration in HeLa-229 cells.

### 2.5 EB and MOMP IgG Quantification by Area Under Dilution Curve

Recombinant MOMP SvA, SvB, and SvC were produced based on the amino acid sequences (NCBI CAX09353.1, ABB51015.1, and WP_024067253.1) with an added N-terminal six histidine tag. Synthetic DNA constructs were codon optimized for expression in *Escherichia coli*, followed by insertion into the pJexpress 411 vector (Atum). Purification was done essentially as described elsewhere (Olsen *et al*., 2010). Briefly, expression was induced by IPTG in *Eschericia coli* BL-21 (DE3) cells transformed with the synthetic DNA constructs. Inclusion bodies were isolated and extracts were loaded on a HisTrap column (GE Healthcare), followed by anion exchange chromatography on a HiTrap Q HP column and dialysis to 20mM tris-HCl, pH 8.0. Protein concentrations were determined by the bicinchoninic acid protein assay.

For EB-specific IgG assays, test and positive control serum were diluted 1:10,000, 1:1000, 1:100 and 1:10. For MOMP-specific IgG assays, test and positive control serum were diluted 1:10,000, 1:500, 1:50 and 1:10. 45μl of diluted serum was added to each well and incubated at room temperature for two hours. Plates were then washed four times. Anti-human IgG HRP antibody (Anti-Human IgG-Peroxidase from Sigma) was diluted 1:30,000 in blocking buffer, and 100ul added to each well. Following incubation at room temperature for one hour, plates were washed four times. 100μl 1-Step Ultra TMB-ELISA Substrate Solution was added per well and the plate developed for 30 minutes in the dark. 100μl 1M sulphuric acid was added per well to stop the reaction. Plates were read at 450nm and 630nm. The absorbance at 630nm was subtracted from the absorbance at 450nm to give a background-adjusted absorbance. For each plate, absorbances were normalised relative to a positive control, fit to a 4-parameter logistic curve using the drc package in R, and the area under the curve calculated to give a measure of antigen-specific IgG quantity.

### 2.6 MOMP-Peptide Specific IgG and Validation of MOMP IgG Response

Synthetic peptides (20-22mers) representing variable domain (VD) VD1, VD2, preVD3, VD3 and VD4 from SvA, B and C were produced by Genecust (Table S1). Maxisorp Plates (Nunc, Denmark) were coated overnight with either recombinant MOMP (1μg/ml) or synthetic peptides (10μg/ml) in carbonate buffer. Plates were washed with PBS containing 0.2% Tween-20 and blocked with 2% SM in PBS. Serum samples were diluted 1:50 in PBS with 1% SM, and as secondary antibody we used anti-human IgG HRP antibody (Agilent P0214) diluted 1:8,000 in PBS + 1% SM. TMB Plus (Kem-En-TEC) was used for detection and plates were read at 450nm with subtraction of the absorbance value measured at 620nm. The MOMP peptide analysis was performed for 37 samples with a positive response to either recombinant SvA, SvB or SvC MOMP.

### 2.7 *C. trachomatis* Neutralisation

For 91 out of 93 samples the neutralization assay was performed essentially as described (Byrne 1993, Olsen 2021). Briefly, Syrian golden hamster kidney (HaK) cells (ATCC CCL-15) were seeded in 96-well flat-bottom microtiter plates. Serum samples were heat-inactivated at 56°C for 30 minutes and diluted in SPG buffer to create two-fold dilution series starting from 20-fold dilution. The diluted *C. trachomatis* SvB stock was mixed 1:1 with the diluted plasma samples and incubated for 45 minutes at 37°C, 5% CO2. The mixture was used for infection of HaK cells for two hours at 35°C and thereafter the cells were incubated for 24 hours. Chlamydial inclusions were visualized by staining with polyclonal rabbit anti-recombinant CT043 serum, followed by Alexa Fluor 488–conjugated goat anti-rabbit immunoglobulin (Life Technologies). Cells were stained with propidium iodide (Invitrogen) and inclusion forming units (IFUs) were counted by an ImageXpress Pico automated cell imaging system (Molecular Devices, San Jose, CA) as previously described (Olsen *et al*., 2021). A positive and negative reference pool from rabbits immunized with Hirep1 (Olsen *et al*., 2015) were included on all plates. Percent specific neutralisation was calculated as [(No sample control IFU – sample IFU) / No sample control IFU] × 100 for each dilution, and the serum dilution giving a 50% reduction in IFU was named reciprocal 50% neutralisation titer (NT50). Reciprocal NT50 values were calculated based on a five parameter logistic curve using the package ‘drc’ in R. For samples where no titer could be calculated, the samples were assigned a NT50 titer of 10 (half the value of the lowest dilution).

### 2.8 Fc-Receptor Dependent Phagocytosis

The phagocytosis assay has been previously described (Grasse *et al*., 2018), with the modification that *C. trachomatis* was not prelabeled with carboxyfluorescein diacetate succinimidyl ester in this study. Briefly, PLB-985 cells, a human myeloid leukaemia line, were cultured in RPMI 1640 with 1% L-Glutamine, 1% HEPES, 1% pyruvate, 1% Non-Essential Amino Acids Solution, 10 μg/ml gentamicin, and 10% heat-inactivated FBS at 37°C and 5% CO2. PLB-985 cells were stimulated with 100mM N,N-Dimethylformamide (DMF; Sigma-Aldrich) for five days to induce differentiation into a neutrophil-like phenotype. Serum samples were diluted 1:100 and *C. trachomatis* SvB bacteria were diluted in the PLB-985 culture medium, mixed in a 1:1 ratio, then incubated for 40 minutes at 37°C on a rocker table. 40μl of the serum/bacteria suspension was then mixed with 100μl of the DMF stimulated PLB-985 cells (2 × 10^6^ cells/ml). Following incubation for four hours at 37°C on a rocker table, cells were washed with PBS, then stained with eBioscience Fixable Viability Dye eFluor 520. After 15 minutes, cells were washed with FACS buffer (PBS with 2% FBS, 0.1% sodium azide, 1mM EDTA) and fixed using BD Cytofix. Thereafter, cells were incubated with monoclonal mouse anti-*C. trachomatis* LPS (Abnova) in PBS followed by incubation with goat-anti-mouse IgG conjugated with AlexaFluor 647. Samples were measured using a BD FACSCanto. Live singlet PLB-985 cells were gated and the *C. trachomatis* SvB signal measured in the APC channel. The analysis was performed for a representative subset of 26 samples.

### 2.9 Serology Data Analysis

Statistical analysis was carried out in the R statistical software environment. Correlation between antibody features was assessed by calculating Spearman’s correlation coefficient for complete observations. Distance was calculated using the as.dist function in R (distance = as.dist(1-Spearman’s rank correlation co-efficient)). Unsupervised hierarchical cluster analysis was then performed on the distance matrix using the hclust function in R. A test of the correlation coefficient being zero was carried out using the cor.test function.

To find whether any individual antibody features were associated with susceptibility, each antibody feature was centered and scaled, and a logistic regression model with age, sex and ethnicity as covariates used. To carry out multivariate analyses, missing data was imputed using the k-nearest neighbors (kNN) method in the preprocessing function of the caret package (Kuhn 2008). Briefly, kNN imputation identifies the samples most similar to the sample with missing data (the nearest neighbours), and averages the value from these samples. Data was once again centered and scaled. Principal component analysis was carried out using the prcomp function in R. A partial least squares classification model was built using the caret package, using receiver operating characteristic (ROC) as a metric to choose the optimal model. In this case a model using two components (linear combinations of the original features) was optimal. The specificity and sensitivity of the model in classifying susceptible individuals was tested using 10-fold cross-validation.

## 2 Results

### 3.1 Infection Susceptibility in the Study Population

Serological assays were carried out on serum samples from n = 93 children living in 19 family compounds in the Gambia. These children were a subgroup of those recruited to a larger longitudinal study (n = 345, Faal 2006). Family compounds in the longitudinal study were visited every two weeks for 28 weeks (15 time points) to test for ocular *C. trachomatis* infection. In addition to the 93 children who underwent serological profiling at baseline, a further 67 children in these 19 compounds were followed up longitudinally for infection (Figure 1a, total of 160 children in these compounds). By a combination of whole genome sequencing and ompA amplicon sequencing, ompA serovar was successfully determined for at least one time point in 47 individuals, of which 40 were infected with serovar B; 5 with serovar A; and two individuals with A and B at separate time points (Figure 1b).

**Figure 1:**
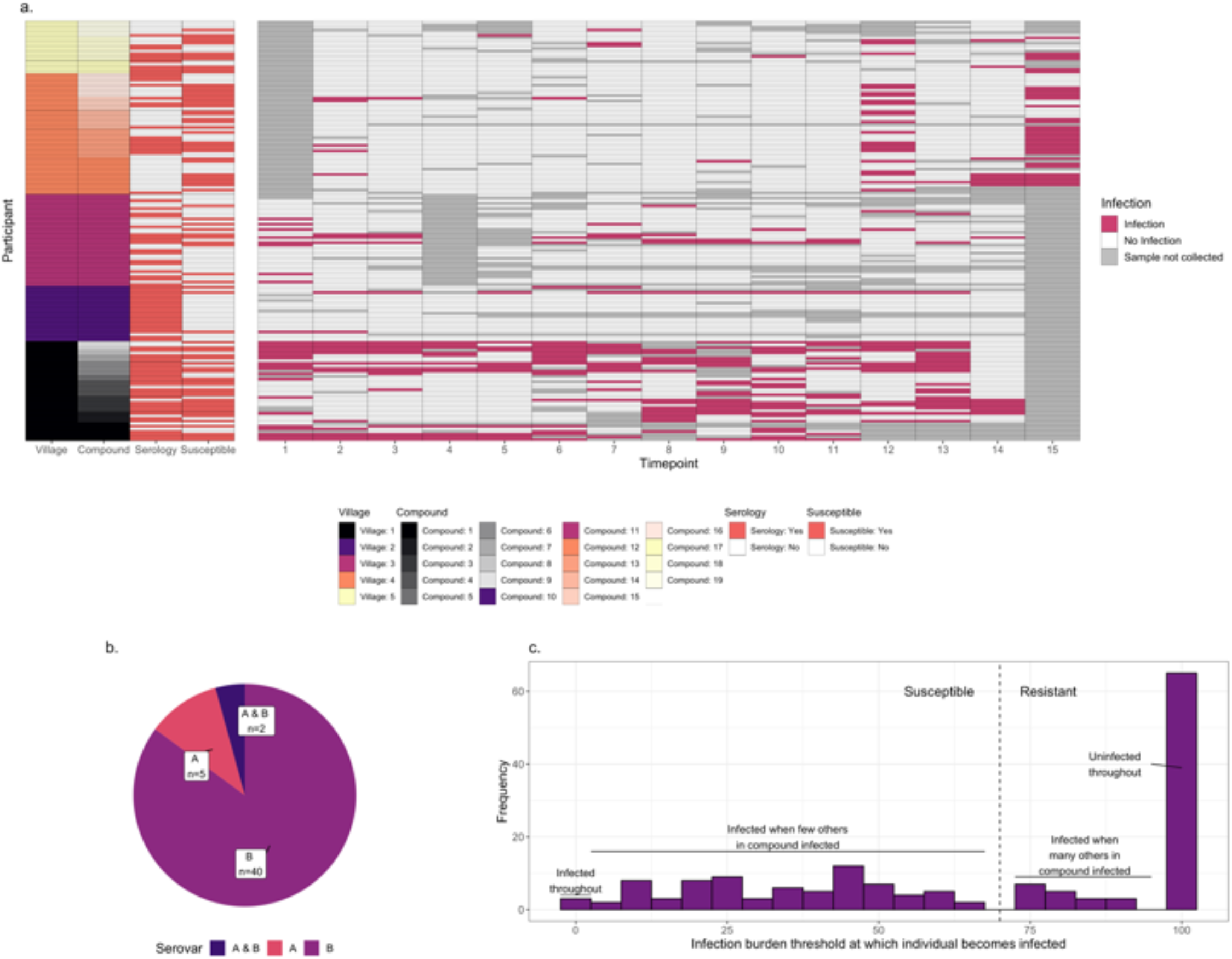
a. *C. trachomatis* infection over fifteen time points (two week intervals) in 160 children from five villages and 19 compounds in the Gambia. Infected time points are highlighted in red, whereas timepoints where a sample was not collected for a particular participant is shown in grey. The characteristics of each participant are annotated on the left, including village, compound, whether they were deemed susceptible (red = yes), and whether serological assays were carried out (red = yes). Assays were carried out on samples collected at time point T1. b. For 51 individuals for which *C. trachomatis* serovar was successfully identified, the proportion which were infected with serovar A, infected with serovar B, or infected with both at different timepoints. c. Distribution of the infection burden (% infected) a child’s compound would need to reach before they themselves became infected. 160 individuals from 19 compounds, of which a subset were sampled for systems serology (93/160) are included.

The proportion of sampled children who were *C. trachomatis* positive varied over time and between compounds. For each of the 160 participants in the 19 compounds from which samples were collected, we examined how the percentage of *C. trachomatis* positive children in their compound varied over time. A logistic regression model was used to predict what threshold the infection burden in their compound needed to reach before each individual became infected (Figure S1). This threshold followed a bimodal distribution (Figure 1c). We therefore considered children who became infected when less than 70% of the other children in their compound were infected to be “susceptible”, and others to be “resistant”.

Of the 93 children that underwent serological profiling at baseline, n = 45 were susceptible and n = 48 resistant. While the children in these two groups were similar in age, female children were more likely to be susceptible (susceptible group 60% female versus resistant group 31% female, p=0.0069, fisher test; Table 1, Table S2).

**Table 1:** Characteristics of “susceptible” and “resistant” participants for the 93 samples on which serological assays were performed.

|  | Susceptible | Resistant |
| --- | --- | --- |
| N | 45 | 48 |
| Age | 7 (IQR 5-10) | 8 (IQR 6-12) |
| % Female | 60 | 31.25 |
| Ethnicity | Mandinka 31%,<br>Manjago 51%,<br>Serahule 7%, Serer 2%, Wolof 9 % | Mandinka 56%,<br>Manjago 15%,<br>Serahule 29 % |

### 3.2 Antigen-Specific Antibody Responses to Different Serovars Correlate Closely With One Another

The serological assays carried out are summarized in Figure 2. Antigen-specific assays measured total IgG specific to *C. trachomatis* EBs and recombinant MOMP (both calculated by area under the dilution curve), and five peptides within or adjacent to the variable domains (VD) of MOMP (serovars A, B and C). Functional assays measured the ability of serum to neutralize infection of HaK cells with serovar B *C. trachomatis*, or to induce Fc-receptor dependent phagocytosis of serovar B *C. trachomatis* by myeloid PLB-985 cells.

**Figure 2:**
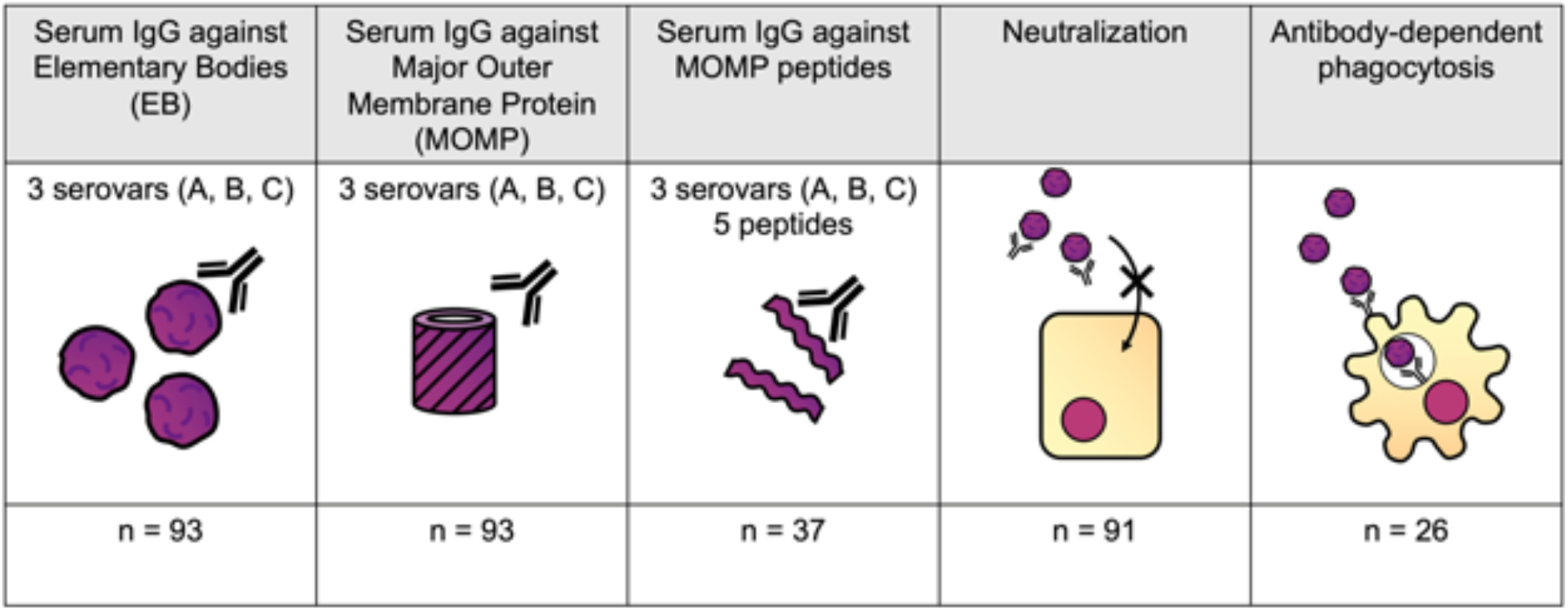
Serological assays performed. Antigen-specific assays measured total IgG specific to *C. trachomatis* EBs, recombinant MOMP, and five peptides within MOMP (serovars A, B and C). Functional assays measured the ability of serum to neutralize infection of HaK cells, or to induce Fc-receptor dependent *C. trachomatis* phagocytosis by myeloid PLB-985 cells.

To elucidate the relationship between different antibody features, Spearman’s rank correlation coefficient was calculated for each pair of features. This correlation matrix was used to calculate a distance matrix and hierarchal clustering carried out. Antibody features clustered by whether the assay measured functional responses, IgG responses to EB and MOMP or IgG responses to MOMP peptides (Figure 3a). Antigen-specific IgG responses to whole EBs and MOMP closely correlated with one another, regardless of the antigen serovar (Figure 3b; Spearman’s rank correlation coefficients ranging from 0.692-0.998, median 0.820). IgG responses to peptides within MOMP were also closely correlated (Spearman’s rank correlation coefficients ranging from 0.160-0.979, median 0.715), although the response to VD1 SvB was a notable outlier, with generally low responses. Fc-dependent phagocytosis correlated more strongly with neutralization than any other feature (Spearman’s rank correlation coefficient of 0.502).

**Figure 3:**
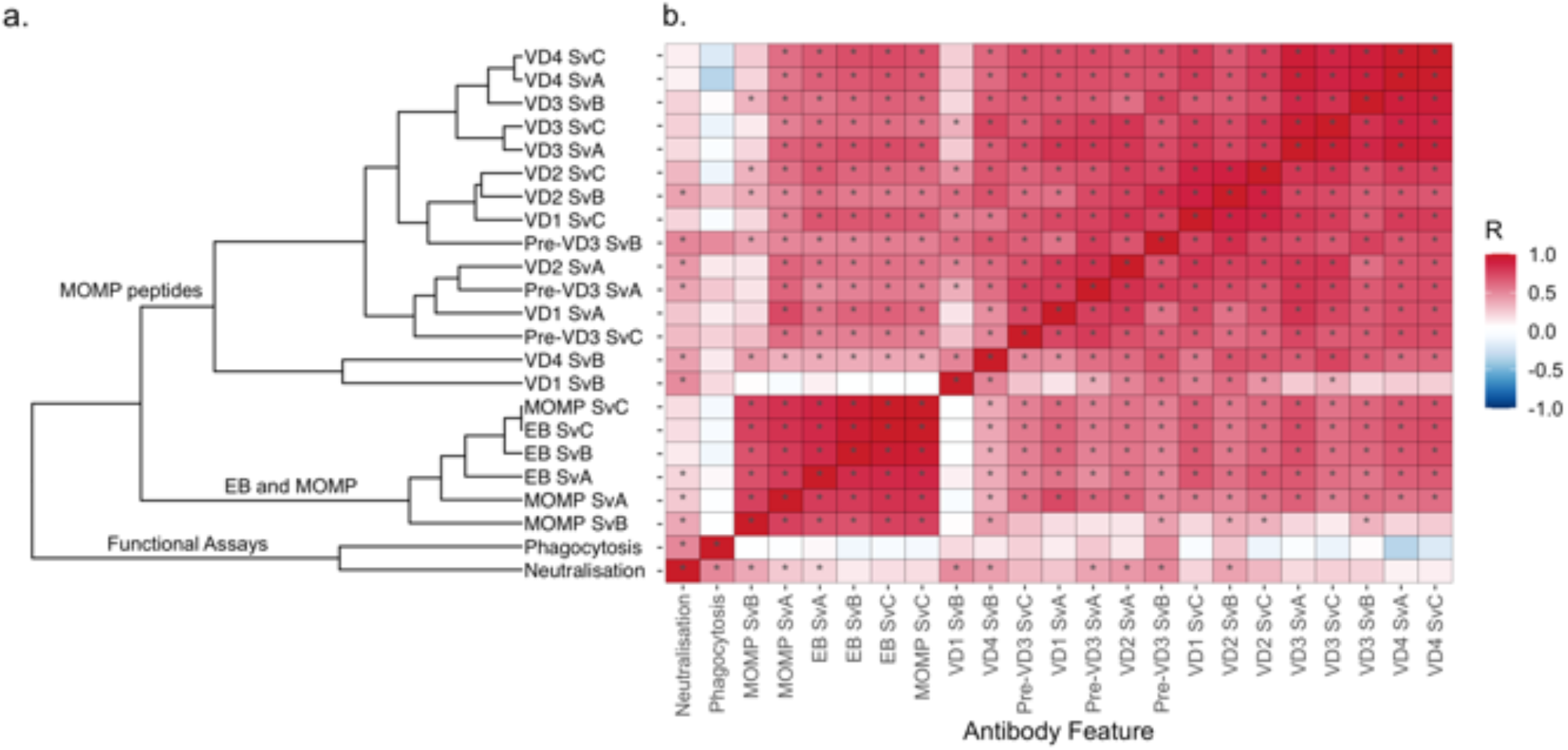
a. Hierarchal clustering of antibody features. Spearman’s rank correlation co-efficient was calculated for each pair of features, and distance calculated using the as.dist function in R (distance = as.dist(1-Spearman’s rank correlation co-efficient)). Unsupervised hierarchical cluster analysis was then performed on the distance matrix. b. Correlation between different antibody features, colored by spearman’s rank correlation coefficient. Features are ordered as in Figure 3a. Significant correlations (p < 0.05) are highlighted by an asterisk.

### 2.3 MOMP-Specific Antibody Responses Were Not Associated with Resistance to Infection

A logistic regression model was used to assess the association of each feature with susceptibility to infection over the following six months, adjusting for age, sex and ethnicity (Figure 4a and Table 2). No features were significantly associated with resistance (p < 0.05), but IgG responses towards MOMP serovar A (p = 0.0022, false discovery rate = 0.052) and neutralization titer (p = 0.013, false discovery rate = 0.15) were higher in susceptible individuals (Figure 4b). The higher levels of IgG against MOMP serovar A in susceptible individuals was validated in an ELISA assay carried out at a different site, and carried out at a single serum dilution rather than by area under dilution curve (Figure 4c; p = 0.0003, logistic regression model).

**Figure 4:**
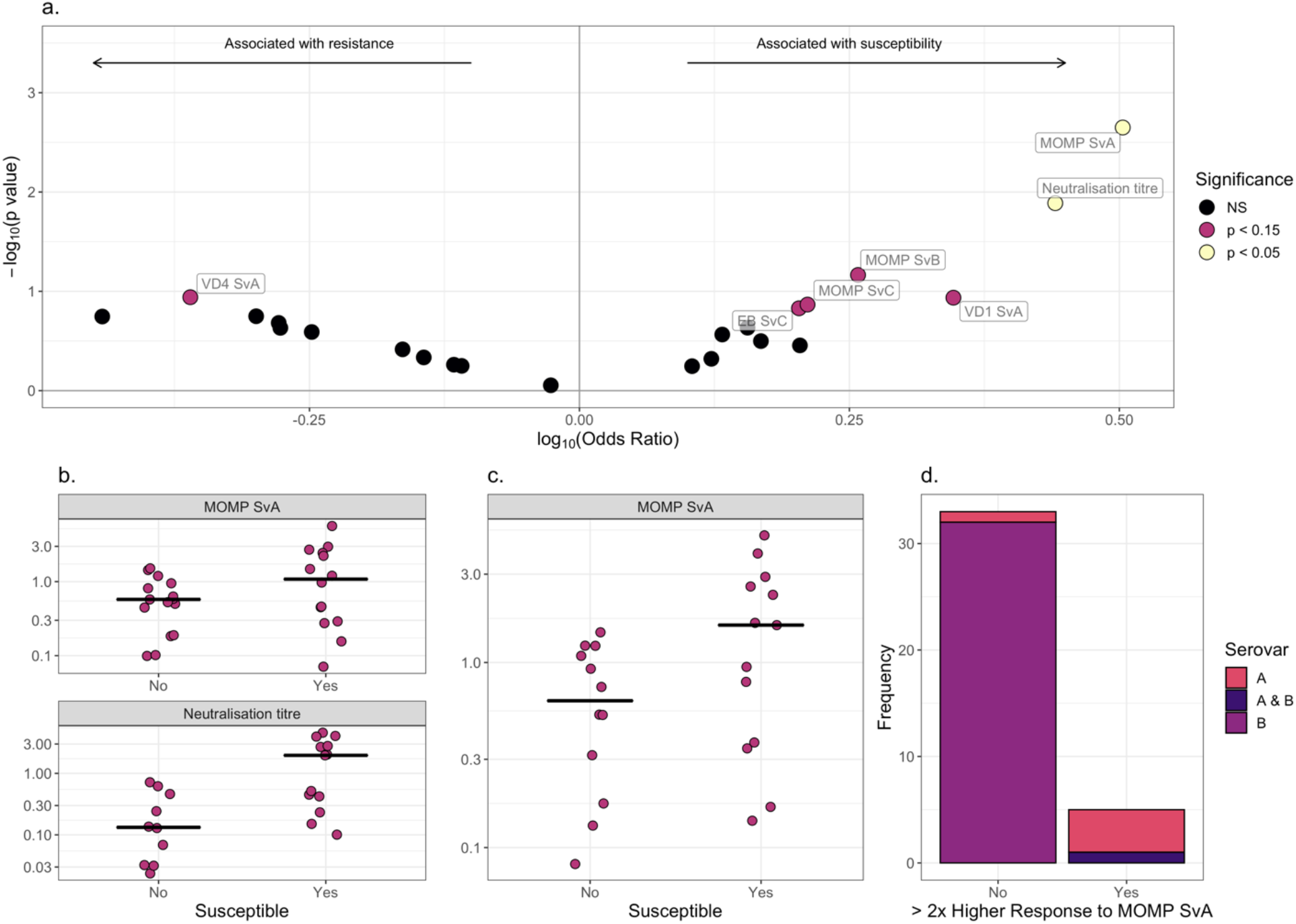
a. Volcano plot for association between different antibody features and susceptibility to infection. Log-transformed odds-ratio is shown on the x axis, with a log_10_(Odds ratio) > 0 indicating association with susceptibility in a logistic regression model where age, sex and ethnicity were included as covariates. On the y axis, features more significantly associated with susceptibility have a greater value of –log_10_(p value). Points are colored by significance level: non-significant (NS) in black, p < 0.15 in pink and p < 0.05 in yellow. b. Comparison of scaled anti-MOMP SvA IgG and Neutralization titer between susceptible and resistant participants. Median is indicated by a horizontal line. Values are shown on a log_10_ transformed axis. c. Comparison of scaled anti-MOMP SvA IgG between susceptible and resistant participants for an ELISA carried out at a different site. Median is indicated by a horizontal line. Values are shown on a log_10_ transformed axis. d. Serovars that participants with high baseline anti-MOMP SvA IgG (>2x response to MOMP SvB IgG) became infected with, versus serovars that participants without high baseline anti-MOMP SvA IgG became infected with.

**Table 2:** Association between different antibody features and susceptibility to infection. Log_10_(Odds ratio) > 0 indicating association with susceptibility in a logistic regression model where age, sex and ethnicity were included as covariates. 95% confidence intervals are indicated.

| Antibody feature | $\text{Log}_{10}(\text{Odds ratio})$<br>(95% confidence intervals) | p value | False discovery rate |
| --- | --- | --- | --- |
| Higher in susceptible individuals |  |  |  |
| MOMP SvA | 0.5 (0.23 to 0.89) | 0.0022 | 0.052 |
| Neutralization titer | 0.44 (0.15 to 0.89) | 0.013 | 0.15 |
| MOMP SvB | 0.26 (-0.00077 to 0.56) | 0.068 | 0.45 |
| VD1 SvA | 0.35 (-0.022 to 0.91) | 0.12 | 0.45 |
| MOMP SvC | 0.21 (-0.03 to 0.53) | 0.14 | 0.45 |
| EB SvC | 0.2 (-0.036 to 0.52) | 0.15 | 0.45 |
| EB SvB | 0.16 (-0.072 to 0.45) | 0.23 | 0.45 |
| EB SvA | 0.13 (-0.09 to 0.39) | 0.27 | 0.45 |
| VD2 SvA | 0.17 (-0.16 to 0.53) | 0.32 | 0.49 |
| Phagocytosis | 0.2 (-0.21 to 0.76) | 0.35 | 0.5 |
| Pre-VD3 SvA | 0.12 (-0.23 to 0.49) | 0.48 | 0.58 |
| Pre-VD3 SvC | 0.1 (-0.26 to 0.49) | 0.57 | 0.59 |
| Higher in resistant individuals |  |  |  |
| VD4 SvA | -0.36 (-0.91 to 0.043) | 0.11 | 0.45 |
| VD2 SvC | -0.44 (-1.2 to 0.068) | 0.18 | 0.45 |
| VD4 SvC | -0.3 (-0.81 to 0.094) | 0.18 | 0.45 |
| VD3 SvB | -0.28 (-0.8 to 0.12) | 0.21 | 0.45 |
| VD3 SvC | -0.28 (-0.83 to 0.12) | 0.23 | 0.45 |
| VD3 SvA | -0.25 (-0.77 to 0.14) | 0.26 | 0.45 |
| VD4 SvB | -0.16 (-0.55 to 0.21) | 0.38 | 0.52 |
| VD1 SvC | -0.14 (-0.57 to 0.23) | 0.46 | 0.58 |
| Pre-VD3 SvB | -0.12 (-0.51 to 0.27) | 0.55 | 0.59 |
| VD2 SvB | -0.11 (-0.5 to 0.26) | 0.56 | 0.59 |
| VD1 SvB | -0.026 (-0.41 to 0.33) | 0.88 | 0.88 |

For 38 individuals both serovar and serology data were available. While IgG responses against MOMP serovar B were on average similar to MOMP serovar A (median 1.07 vs 1.01), six of these individuals had a baseline response against serovar A over twice the magnitude of their response to B. These individuals were significantly more likely to later become infected with serovar A or both A and B (Figure 4d, p = 1 x 10^-5^, fisher-test), suggesting that lack of protection is not due to serovar-specific immunity.

### 2.4 Supervised Machine Learning Is Not Able to Distinguish Between Susceptible and Resistant Participants Based on Antibody Profile

Finally, as no individual antibody features were associated with resistance, we next investigated whether those who were resistant to infection had a broadly distinct antibody profile from those who were susceptible. Following imputation of missing data using k-Nearest Neighbors, principal component analysis based on all features did not reveal any major difference in overall serological profile between groups (Figure 5a). 10-fold cross-validation of a partial least squares model gave a specificity of 71%, sensitivity of 36% and ROC of 0.58 in distinguishing susceptible from resistant individuals (Figure 5b).

**Figure 5:**
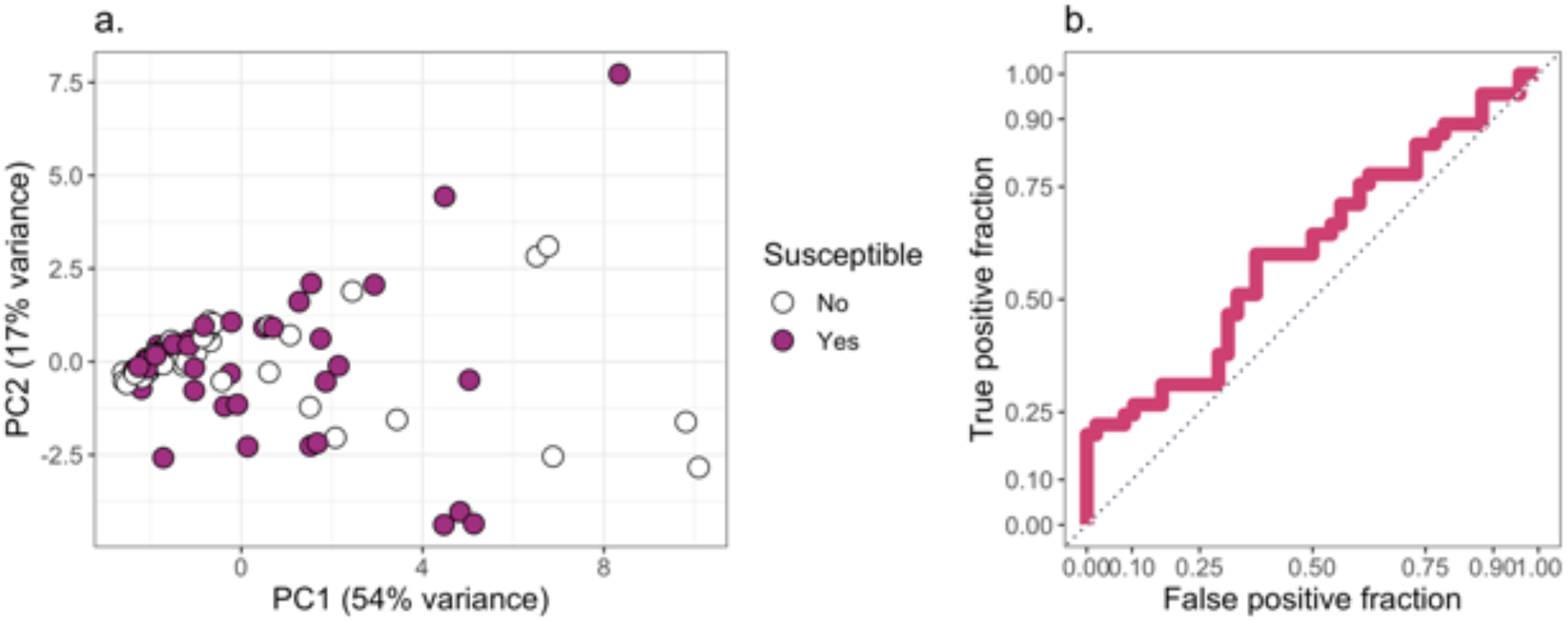
a) Principal components 1 and 2 for a principal component analysis carried out on all features. Missing data was imputed using k-nearest neighbors. Points are colored by whether a participant was classed as resistant or susceptible b) True positive rate against false positive rate for a partial least squares model used to classify participants into “susceptible” or “resistant” categories.

Collectively, our results indicate that neither individual antibody features (p < 0.05) nor overall antibody profile (ROC of 0.58) are associated with protection against ocular *C. trachomatis* infection.

## 3 Discussion

We used a novel definition of susceptibility to identify whether antigen-specific or functional antibody responses were associated with susceptibility to ocular *C. trachomatis* infection. Interestingly, compared with male children, female children became more often infected when the infection burden in their family compound was lower. Although this has not previously been observed for infection, this is consistent with previous studies finding a greater burden of intense trachomatous inflammation, scarring and trichiasis in females, even at a young age (Atik *et al*., 2006; Ramadhani *et al*., 2017; Dolin *et al*., 1998; West *et al*., 1991). This is generally thought to be due to the caring responsibilities of women and female children, resulting in more time in proximity to the age group with the highest infection burden: children aged five and under (Courtright and West, 2004).

Overall, the serological profiles of those who were susceptible and resistant were relatively similar, with a partial least squares model only performing slightly better than chance in classifying participants. Serum MOMP and EB IgG responses to different *C. trachomatis* serovars closely correlated to one another and were higher in those considered susceptible. This is consistent with previous studies finding that those with high levels of ocular *C. trachomatis*-specific IgG were at a higher risk of subsequently developing active trachoma (Bailey *et al*., 1993) and that serum IgG titers to EBs are higher in participants with trachomatous scarring (Holland *et al*., 1993).

One potential explanation for the results observed here is that some unmeasured environmental or innate risk factor is increasing both past infections, which is reflected in high MOMP/EB IgG responses, and further infections after baseline. For example, participants with poor innate immune responses may have had more prolonged infections in the past, resulting in a greater antibody response.

However, we do not rule out the possibility of antibody-dependent enhancement, most famously associated with dengue fever. In secondary dengue virus infections, cross-reactive poorly-neutralizing antibodies against a heterologous serotype can enhance infection of Fc-receptor expressing cells (Diamond and Pierson, 2015). Like dengue, *C. trachomatis* is an intracellular pathogen with multiple circulating serovars, and here antibody-dependent phagocytosis was somewhat higher in susceptible individuals. Serum IgG from women with a recent urogenital *C. trachomatis* infection, or monoclonal antibodies against *C. trachomatis*, can in some cases enhance *in vitro* uptake of EBs, presumably via an Fc receptor on target cells (Ardizzone *et al*., 2021). On the other hand, we found neutralizing antibodies to also be higher in susceptible individuals, and there was no evidence of increased *C. trachomatis* burden in secondary infections.

It is clear that infection-induced anti-MOMP and EB IgG, at least at the levels reached by the children in our cohort, does not protect against re-infection. One possibility is that as the children in our study were relatively young, their IgG may not have yet reached a protective threshold. Another possibility is that the scope of the assays, in focusing on EB and MOMP-specific IgG responses in serum, was too limited. For example, it has previously been found that serum anti-EB IgA titers are higher in healthy controls compared to those with trachomatous scarring (Holland *et al*., 1993), and anti-*C. trachomatis* ocular IgA titers are (non-significantly) higher in those protected from acquiring trachomatous disease (Bailey 1993). Antibody responses to less well characterized antigens could also be protective. For example, PmpD vaccination induces neutralizing, non-serovar specific antibodies, and protects against murine intra-vaginal *C. trachomatis* infection (Crane *et al*., 2006; Paes *et al*., 2016). Protein arrays have also identified a number of potentially protective antibody responses, including those against HtrA, CT043, CT016, Nqr3, and Tarp, which protect against nasal *C. muridarum* challenge in mice (Finco *et al*., 2011), and anti-CT442 responses, which are higher in those without scarring in a trachoma-endemic area (Pickering *et al*., 2017). Furthermore, local ocular responses may have a different effect than serum responses. Finally, both mouse and human studies have shown *C. trachomatis*-specific CD4^+^ IFN-γ responses to be highly important for protection, and therefore humoral responses may play a redundant role in immunocompetent individuals (Morrison *et al*., 2000; Bakshi *et al*., 2018).

This study was limited in that certain assays were not carried out for every participant, and systemic rather than ocular responses were measured. However, as participants were longitudinally monitored for infection following baseline blood sample collection, we were uniquely placed to examine how antibody profile preceding infection relates to susceptibility. We find that infection-induced antibody responses towards MOMP do not appear to be protective in this setting, suggesting that natural immunity may be mediated by other mechanisms, or develop at an older age than the participants included in this study.

## 5 Author contributions

AB led the statistical and multivariate analysis, with significant contribution from MJH, IR and JD. AB wrote the first draft, which was reviewed and edited by IR, HP, NF, AH, HJ, PM, MR, AWO, RB, DCWM, FF, JD and MJH. NF, HJ, PM, RB and DCWM were involved in clinical study sample collection. IR, JD and AWO designed the experiments and methodology, and IR and AWO provided reagents. Experiments were carried out by IR (peptide and MOMP ELISA, phagocytosis and neutralization assays) HP (EB and MOMP ELISA), AH and MR (*C. trachomatis* sequencing).

## 7 Acknowledgements

We would like to thank the study participants, as well as the trachoma field and technical teams, including Isatou Sarr, Esther Aryee, Omar Camara and Omar Manneh.

